# Aberrant light sensing and motility in the green alga *Chlamydomonas priscuii* from the ice-covered Antarctic Lake Bonney

**DOI:** 10.1101/2023.02.02.526531

**Authors:** Mackenzie Poirier, Pomona Osmers, Kieran Wilkins, Rachael M. Morgan-Kiss, Marina Cvetkovska

**Author notes:** **Corresponding author:** Marina Cvetkovska, Department of Biology, University of Ottawa, 30 Marie-Curie Pr., Ottawa, ON, Canada, K1N 6N5.

## Abstract

The Antarctic green alga *Chlamydomonas priscuii* is an obligate psychrophile and an emerging model for photosynthetic adaptation to extreme conditions. Endemic to the ice-covered Lake Bonney, this alga thrives at highly unusual light conditions characterized by very low light irradiance (<15 μmol m^-2^ s^-1^), a narrow wavelength spectrum enriched in blue light, and an extreme photoperiod. Genome sequencing of *C. priscuii* exposed an unusually large genome, with hundreds of highly similar gene duplicates and expanded gene families, some of which could be aiding its survival in extreme conditions. In contrast to the described expansion in the genetic repertoire in *C. priscuii*, here we suggest that the gene family encoding for photoreceptors is reduced when compared to related green algae. This alga also possesses a very small eyespot and exhibits an aberrant phototactic response, compared to the model *Chlamydomonas reinhardtii*. We also investigated the genome and behaviour of the closely related psychrophilic alga *Chlamydomonas* sp. ICE-MDV, that is found throughout the photic zone of Lake Bonney and is naturally exposed to higher light levels. Our analyses revealed a photoreceptor gene family and a robust phototactic response similar to those in the model *Chlamydomonas reinhardtii*. These results suggest that the aberrant phototactic response in *C. priscuii* is a result of life under extreme shading rather than a common feature of all psychrophilic algae. We discuss the implications of these results on the evolution and survival of shade adapted polar algae.

## Introduction

Light provides energy and information that regulates many cellular processes in plants and algae. Motile green algae have sensitive mechanisms for light detection and can induce movement across a light gradient, either towards (positive phototaxis) or away from a light source (negative phototaxis) (Böhm & Kreimer, 2020). Phototaxis is regulated by a specialized organelle called an eyespot that allows for the precise detection of light intensity and direction (Ueki *et al*., 2016; Böhm & Kreimer, 2020). Photoreceptors are critical components of the light-sensing apparatus that controls phototaxis and a plethora of other processes including photosynthesis, circadian rhythms, and gametogenesis (Trippens *et al*., 2012; Petroutsos *et al*., 2016; Müller *et al*., 2017; Rredhi *et al*., 2021). Phototaxis has evolved independently multiple times in diverse microbial lineages including cyanobacteria, algae, and protists (Jékely, 2009; Gavelis *et al*., 2017), suggesting that light-directed movement confers an evolutionary advantage that allows free-swimming microbes to avoid stress from either insufficient or excess light.

Many important insights on light sensing come from model species, such as *Chlamydomonas reinhardtii* (Kianianmomeni & Hallmann, 2014), but green algae are found in diverse habitats. Many environments are populated by species thriving under environmental regimes that are untenable for growth of most model algae. Lake Bonney of the McMurdo Dry Valleys in Antarctica is one such environment. Microalgal communities in this lake are challenged with perpetual low temperatures, extreme shading under a perennial ice cover, prolonged periods of darkness during the polar winter, nutrient deficiencies, supersaturated oxygen levels, and high salinity (Morgan-Kiss *et al*., 2006; Patriarche *et al*., 2021). The ice cover prevents wind-driven mixing and environmental inputs, making this lake an unusually stable and highly stratified environment, often termed a “natural laboratory” for the study of extremophilic biology (Patriarche *et al*., 2021). Lake Bonney is a home to a diverse algal community, including one of the best studied polar chlorophytes *Chlamydomonas priscuii*, recently re-named from *Chlamydomonas* sp. UWO241 (Stahl-Rommel *et al*., 2021).

Chlorophytes dominate the phytoplankton communities of Lake Bonney. *C. priscuii* has only been detected in the deep photic zone at 17 meters below the surface of the ice (Neale & Priscu, 1995) while a second chlorophyte, *Chlamydomonas* sp. ICE-MDV is found throughout the photic zone and is the dominant chlorophyte within the shallower, under-ice layers (Bielewicz *et al*., 2011). In its natural environment *C. priscuii* is exposed to year-round low temperatures (∼4°C), hypersalinity (700 mM NaCl), low light irradiance (<15 μmol m^-2^ s^-1^) with a narrow spectral range (450-550 nm), and long periods of darkness during the polar night (Neale & Priscu, 1995). *C. priscuii* is an obligate cold extremophile (psychrophile) that experiences heat shock and cell death at temperature >18°C (Possmayer *et al*., 2011; Cvetkovska *et al*., 2022). Under lab conditions, *C. priscuii* is present as either biflagellate, highly motile single cells or as nonmotile, multi-celled palmelloids. Pocock *et al*. (2004) observed a very small eyespot composed of a single layer of carotenoid-rich globules. Positive phototaxis was only possible at higher temperatures (25°C) but not at those closer to its natural environment (7°C) (Pocock *et al*., 2004). In native phytoplankton communities, Priscu and Neale (1995) observed that while there is no evidence of diel migration in the water column, shallow phytoplankton populations exhibited positive phototaxis, while deeper communities (12 m and 20 m sampling depths) did not. The spatial distribution of different *Chlamydomonas* populations within the water column of Lake Bonney, combined with recent advances in the study of *C. priscuii*, including the sequencing of its genome (Zhang *et al*., 2021), prompted us to further investigate its phototactic response.

## Materials and Methods

### Strains and Growth Conditions

*Chlamydomonas priscuii* (previously UWO241, CCMP1619) was originally isolated in early 1990s from the deep photic zone (17 m sampling depth) of the east lobe of Lake Bonney, Antarctica (Neale & Priscu, 1995). *Chlamydomonas* sp. ICE-MDV was originally isolated in 2014 from an enrichment culture of the east lobe of Lake Bonney (Li *et al*., 2016). *Chlamydomonas reinhardtii* (CC-1690) was obtained from the Chlamydomonas Resource Center. All cultures were grown axenically in Bold’s Basal Medium (BBM) supplemented with 700 mM NaCl at 4°C (*C. priscuii*), 70 mM NaCl at 4°C (ICE-MDV), or 0.43 mM NaCl at 24°C (*C. reinhardtii*). All cultures were grown in 500 mL Erlenmeyer flasks continuously aerated with ambient air filtered through a 0.2 μm filter and under continuous light (40 μmol m^-2^ s^-1^) provided by full spectrum LED light bulbs. Light intensity was measured with a quantum sensor attached to a radiometer (Model LI-189; Li-COR). Cell growth was monitored as change in optical density at 750 nm and cell density was measured using a Countess II FL Automated Cell Counter (ThermoFisher Scientific). To ensure that the cultures were viable at the time of the experiment, cell death was measured by labelling with the fluorescent dye SYTOX Green (ThermoFisher Scientific) as described previously (Possmayer *et al*., 2011). To exclude cell mortality as the reason behind the lack of phototaxis, we ensured close to 100% cell viability in each experiment (data not shown). Unless otherwise specified, actively growing cultures in the mid-log phase were used in all experiments. Images of algal cells were taken using a Zeiss Axiophot Microscope (Carl Zeiss AG) on a wet mount slide.

### Identification of photoreceptors genes in green algal genomes

The *C. priscuii* genome (Zhang *et al*., 2021; BUSCO score of 85%) and transcriptome (Cvetkovska *et al*., 2022) were recently sequenced. These datasets were screened for the presence of photoreceptor genes using previously identified sequences from *C. reinhardtii* and conserved photoreceptor domains (Greiner *et al*., 2017) obtained from Phytozome (v6.1) as queries (Merchant *et al*., 2007; Craig *et al*., 2022). Photoreceptor genes in *C. priscuii* were identified through a tBLASTn search (e-value<e^-10^, bit-score>100) and manually inspected for redundant sequences and to ensure correct gene structure annotation. Conserved domains typical for photoreceptors were identified in the *C. priscuii* genome using Pfam (Mistry *et al*., 2021) and NCBI Conserved Domain Database (Lu *et al*., 2020). The genomes of closely related species from the order Chlamydomonadales were obtained from PhycoCosm (Grigoriev *et al*., 2021) and included: *Chlamydomonas eustigma* NIES-2499 (Hirooka *et al*., 2017), *Chlamydomonas incerta* SAG7.73, *Chlamydomonas schloesseri* CCAP 11/173, *Edaphoclamys debaryana* CCAP 11/70 (Craig *et al*., 2021), *Dunalella salina* CCAP19/18 (Polle *et al*., 2017), *Gonium pectorale* NIES-2863 (Hanschen *et al*., 2016), and *Volvox carteri* v.2.1 (Prochnik *et al*., 2010). The genome of the only other psychrophilic Chlamydomonadalean alga available, *Chlamydomonas* sp. ICE-L (Zhang *et al*., 2020) was obtained from GenBank. Photoreceptor genes identified in genome of ICE-L were identical at the nucleotide level as those found in the genome of ICE-MDV (Raymond & Morgan-Kiss, 2013), suggesting that these strains belong to the same species. Multiple sequence alignments were performed using ClustalW (Sievers *et al*., 2011) implemented through Geneious Prime (Biomatters Ltd, Auckland, New Zealand).

### Phototaxis dish assay

Phototaxis dish assay was preformed according to the protocol by Ueki & Wakabayashi (2017) and Ueki *et al*. (2022). Cells from exponentially growing cultures (∼6x10^6^ cells/mL) were resuspended in phototaxis buffer (5 mM HEPES, 0.2 mM EGTA, 1mM KCl, 0.3 mM CaCl; pH 7.2). In all cases, the phototaxis buffer was supplemented with NaCl (0.43 mM, 70 mM, 700 mM) to match the growth conditions for each species. The algae were incubated for 30 minutes under dim red LED light (624 nm; Cree, Inc.) at the growth temperature to promote motility. Phototaxis was observed in a petri dish (35 mm diameter, 10 mm thickness) placed in a dark chamber and illuminated with a unilateral blue (470 nm; Cree, Inc) or green (525 nm; Broadcam Limited) LED light at 1 or 10 μmol m^-2^ s^-1^ for 5 minutes (*C. reinhardtii*) or 15 minutes (*C. priscuii* and ICE-MDV). ROS and their quenchers were previously shown to regulate the phototactic sign in *C. reinhardtii* (Wakabayashi *et al*., 2011). We used H_2_O_2_ (12.23 mM) to induce positive phototaxis and dimethylurea (DMTU; 75 mM) as quencher of H_2_O_2_ to induce negative phototaxis. Unless otherwise specified, all experiments were performed at a temperature corresponding to the growth conditions for each species (24°C for *C. reinhardtii*; 4°C for *C. priscuii* and ICE-MDV). The plates were photographed before and after incubation, and the resulting images were used to compare cell movement. All assays were completed at a minimum of three biological replicates.

The direction and strength of the phototactic response was quantified by determining pixel intensity in the images using ImageJ (Schneider *et al*., 2012) according to a previously described protocol (Ueki *et al*., 2022) with modifications. In brief, images of the same plate before and after the application of unidirectional light were converted to grayscale and color inverted. The ‘before’ image was subtracted from the ‘after’ image. The pixel density of the entire dish (total density) and the half of the dish closest to the light (phototactic cell density) were measured. The phototactic index was calculated as (phototactic cell density)/(total density). A phototactic index of 1 represents a strong positive phototactic response (movement towards the light), a phototactic index of 0 represents a strong negative phototactic response (movement away from the light), and a phototactic index of 0.5 represents no phototactic response (no directional movement in response to light).

### Photoshock assay

The photoshock response was observed according to the protocol described in Ueki & Wakabayashi (2017) and Ueki *et al*., (2022) on a Zeiss ApoTome microscope equipped with a camera (Carl Zeiss AG). Cells were resuspended in phototaxis buffer at a concentration of ∼1x10^6^ cells/mL and incubated in dim red LED light (624 nm) at their growth temperature as described above. To observe non-directional motility, cells were observed at 60 μmol m^-2^ s^-1^ white light. To observe photoshock, cells were observed at dim red light and shocked with a rapid flash of bright white light using a Speedlight 270EX II external flash (Canon Inc.). Videos were created by taking a 15-second series at 62 frames/s and processed by Zen Pro (Carl Zeiss AG). ImageJ was used for video quantification and statistical analysis were performed in RStudio. All assays were completed at a minimum of three biological replicates, with at least 9 fields of view analyzed for each species.

## Results and Discussion

### The psychrophile *C. priscuii* has a reduced repertoire of photoreceptor genes

Screening the nuclear genome of *C. priscuii* revealed only eight (8) full-length photoreceptor genes, a reduced complement compared to most of its green algal relatives (Figure 1a). These genes contained all conserved domains typical for photoreceptors, suggesting functional proteins (Figure 1b). In contrast, the genome of *C. reinhardtii* encodes for at least 15 photoreceptor genes (Greiner *et al*., 2017), which are conserved within the Chlamydomonadales and most species examined within this group encode 14-16 full-length genes. This includes *Chlamydomonas* sp. ICE-L, an Antarctic sea ice alga and a strain of ICE-MDV (Figure 1a; Table S1).

**Figure 1:**
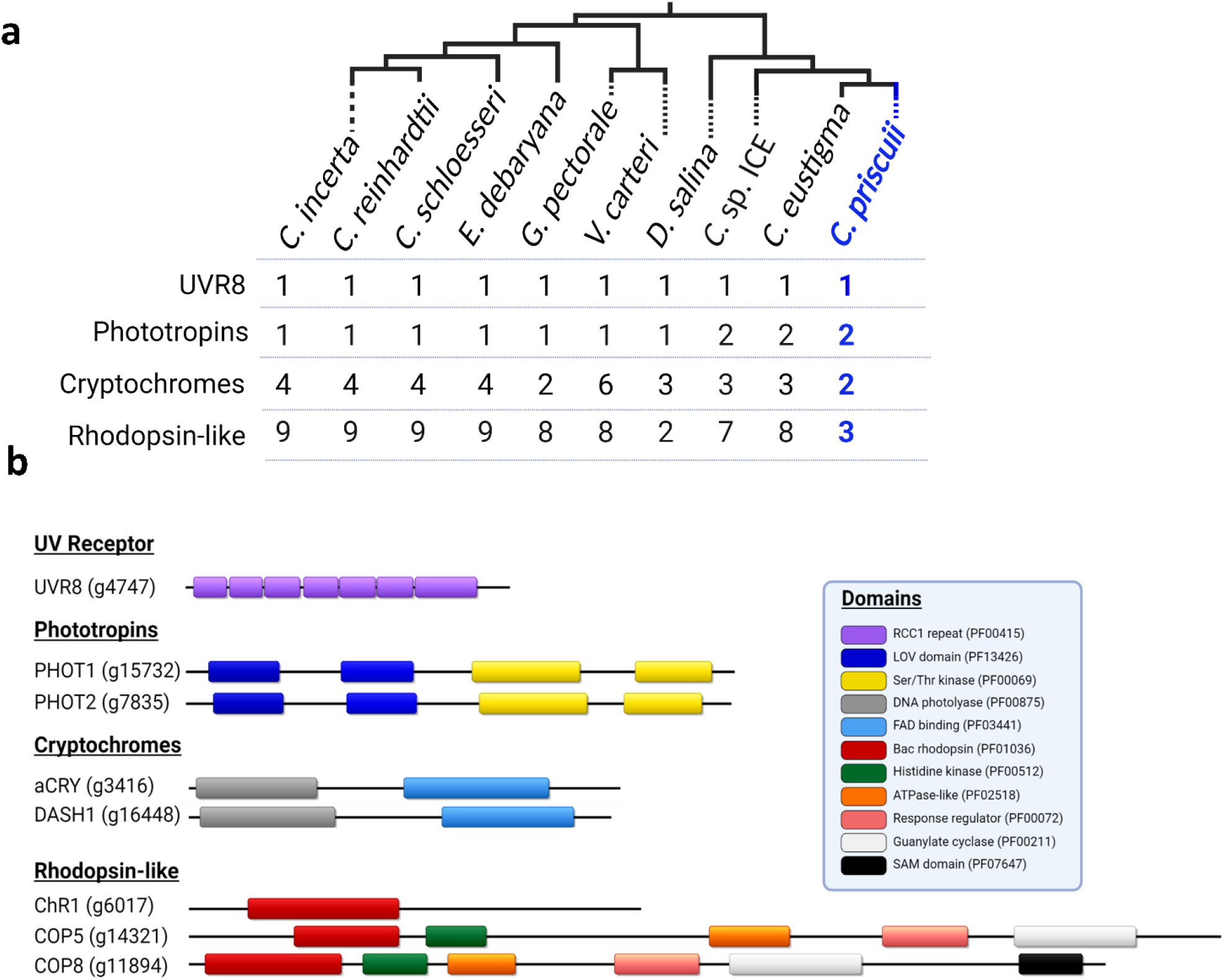
The number of photoreceptor genes and their predicted domain structure in *C. priscuii*. (**a**) Tree of various Chlamydomonadales and the number of full-length photoreceptor genes detected in their nuclear genomes; branching order is based on previous phylogenetic analyses (Possmayer *et al*., 2016; Zhang *et al*., 2020, 2021) and the position of *C. priscuii* is highlighted in blue. (**b**) Overview of the predicted domain structure of the *C. priscuii* photoreceptors, labeled with their gene locus according to Zhang *et al*. (2021). All domains are identified according to their Pfam Family ID. Genome completeness by BUSCO for *C. priscuii* is reported as 85%.

Our results suggest that *C. priscuii* encodes a single channelrhodopsin gene (ChR1), in contrast to two genes in the *C. reinhardtii* genome. The ChR photoreceptors are light-gated cation channels (Berthold *et al*., 2008), and are key determinants of phototactic responses in *C. reinhardtii* (Trippens *et al*., 2012; Greiner *et al*., 2017; Böhm *et al*., 2019; Wakabayashi *et al*., 2021; Baidukova *et al*., 2022). The sites for retinal binding, which are involved in light sensing and protein conformational change (Kato *et al*., 2012) are conserved in *C. priscuii*, suggesting a functional photoreceptor (Figure S1). Böhm *et al*. (2019) proposed that ChR hyperphosphorylation is an important component of phototactic signalling in *C. reinhardtii* but phosphosites are poorly conserved outside of the Reinhardtinia clade with only three predicted phosphosites in the *C. priscuii* ChR compared to twelve in ChR1 in *C. reinhardtii* (Figure S1). Furthermore, none of the conserved phosphosites were in the C-terminal region of the *C. priscuii* ChR, which is of key importance for light-induced ChR1 hyperphosphorylation in *C. reinhardtii* (Böhm *et al*., 2019). Whether these features in the sequence of the gene affect the function and regulation of ChR in *C. priscuii* remains to be experimentally examined.

The gene encoding plant-like cryptochrome (pCRY), that entrains the algal circadian clock in *C. reinhardtii* (Müller *et al*., 2017) was not detected in the *C. priscuii* genome despite being present in all other green algal genomes examined here. The circadian rhythm in *C. priscuii* has not been examined yet, but in its natural environment this alga has a very unusual photoperiod with months-long periods light and dark (Lizotte *et al*., 1996). It appears that *C. priscuii* retains the animal-like cryptochrome (aCRY) and one copy of the phylogenetically conserved *Drosophila, Arabidopsis, Synechocystis, Homo* (DASH) cryptochrome (CRY-DASH), both of which regulate the transcription of genes involved in photosynthesis, chlorophyll biosynthesis, and maintenance of efficient photoautotrophic growth (Beel *et al*., 2012; Rredhi *et al*., 2021). We could also detect only two histidine-kinase rhodopsin genes (COP5 and COP8) in the genome of *C. priscuii*. This in contrast to at least six to eight genes in other green algae (Table S1), including *C. reinhardtii* (COP5-12). The function of histidine-kinase rhodopsin photoreceptors is not well understood in algae (Luck *et al*., 2012; Luck & Hegemann, 2017).

We detected two blue-light receptor phototropins genes (PHOT1 and PHOT2) in the *C. priscuii* genome. Both PHOT genes have the conserved Light-Oxygen-Voltage (LOV) and Ser/Thr kinase domains (Figure 1b) important for blue-light sensitivity and signal transduction (Nakasone *et al*., 2018). ICE-L and *C. eustigma* also share this feature. Having two PHOT receptors is typical for land plants (Li *et al*., 2015), but most unicellular green algae examined to date, including *C. reinhardtii* and its close relatives (Figure 1a) encode a single PHOT receptor. The *C. priscuii* nuclear genome has a high degree of gene duplications (highest of any chlorophyte studied to date), particularly for genes involved in light harvesting and photosynthesis (Zhang *et al*., 2021). Gene duplication is increasingly being viewed as a means of adapting to harsh conditions (Qian & Zhang, 2014). It was hypothesised that *C. priscuii* has retained genes important for life in its cold and shaded environment (Zhang *et al*., 2021). PHOT is involved in the induction of nonphotochemical quenching at high light intensities and is hypothesized to aid in photoprotection (Petroutsos *et al*., 2016).

### The Antarctic *C. priscuii* is motile but exhibits weak photobehaviours

We examined the ability of *C. priscuii* to move in response to light signals and compared it to that of the well-studied responses in the model *C. reinhardtii*. We also tested the phototactic behaviour of its close relative ICE-MDV, isolated from the shallow photic zone in Lake Bonney (Li *et al*., 2016). Natural PAR levels experienced by ICE-MDV (∼50 μmol m^-2^ s^-1^) are approximately fivefold higher compared to that of *C. priscuii* (<15 μmol m^-2^ s^-1^) (Kong *et al*., 2012) allowing for a direct comparison between two closely related psychrophiles from the same environment but adapted to different light conditions. Both psychrophiles exhibited small eyespots, compared to that observed in *C. reinhardtii* (Figure 2a). All algal species had two flagella (Figure 2a) and we confirmed motility under non-directional white light (Movie S1-S3); however, we show that the two Antarctic species swim slower than the mesophilic *C. reinhardtii* (Figure 2b).

**Figure 2:**
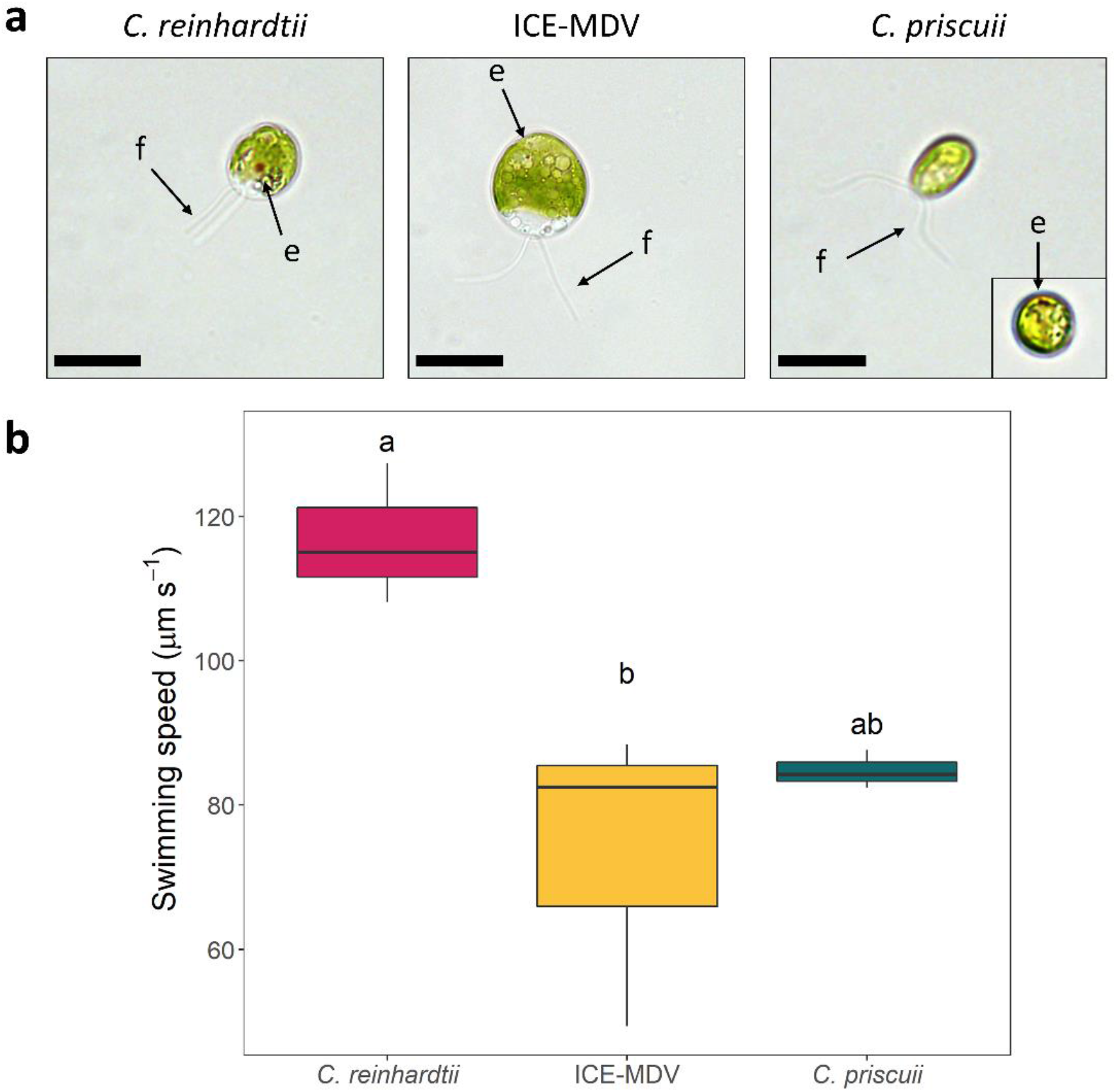
(**a**) Morphology of *C. reinhardtii*, ICE-MDV, and *C. priscuii*. Algae observed under brightfield microscopy exhibit a visible eyespot (e) and two flagella (f), as indicated by black arrows. The image in the inset is a single *C. priscuii* cell where the small eyespot is visible. Scale bar = 10 μm. (**b**) Average swimming speeds of three Chlamydomonadales species. Statistical significance was determined with a one-way ANOVA (p = 0.018) with Tukey’s post hoc test. Statistically different treatments are indicated by different letters.

Using a dish phototaxis assay, we demonstrated that *C. priscuii* exhibits a weaker photobehaviour compared to both ICE-MDV and *C. reinhardtii*. We first exposed algal cultures to unidirectional green light (λ = 525 nm) that regulates phototaxis and has minimal impact on photosynthesis (Ueki & Wakabayashi, 2017). This wavelength induced prominent and rapid negative phototaxis in *C. reinhardtii* after 5 min, even at very low light intensity (1 μmol m^-2^ s^-1^) (Figure 3a, 3b). To account for the slower swimming speeds, we exposed the Antarctic species to this treatment for 15 minutes. We observed strong and consistent positive phototaxis in ICE-MDV after 15-minute exposure to green light even at very low light intensity (1 μmol m^-2^ s^-1^) but the same treatment induced a much weaker and inconsistent phototaxis or no phototactic response in *C. priscuii* (Figure 3a, 3b). We also tested for phototaxis under blue light (λ = 470 nm), the predominant wavelength in the depths of Lake Bonney. Once again, we observed weak or no phototaxis in *C. priscuii*. ICE-MDV exhibited a strong positive phototaxis under blue light, whereas *C. reinhardtii* also moved rapidly but away from the light (Figure S2). These results suggest that *C. priscuii* exhibits a weak and inconsistent phototactic behaviour under environmentally relevant light conditions, compared to a robust response in both *C. reinhardtii* and ICE-MDV.

**Figure 3:**
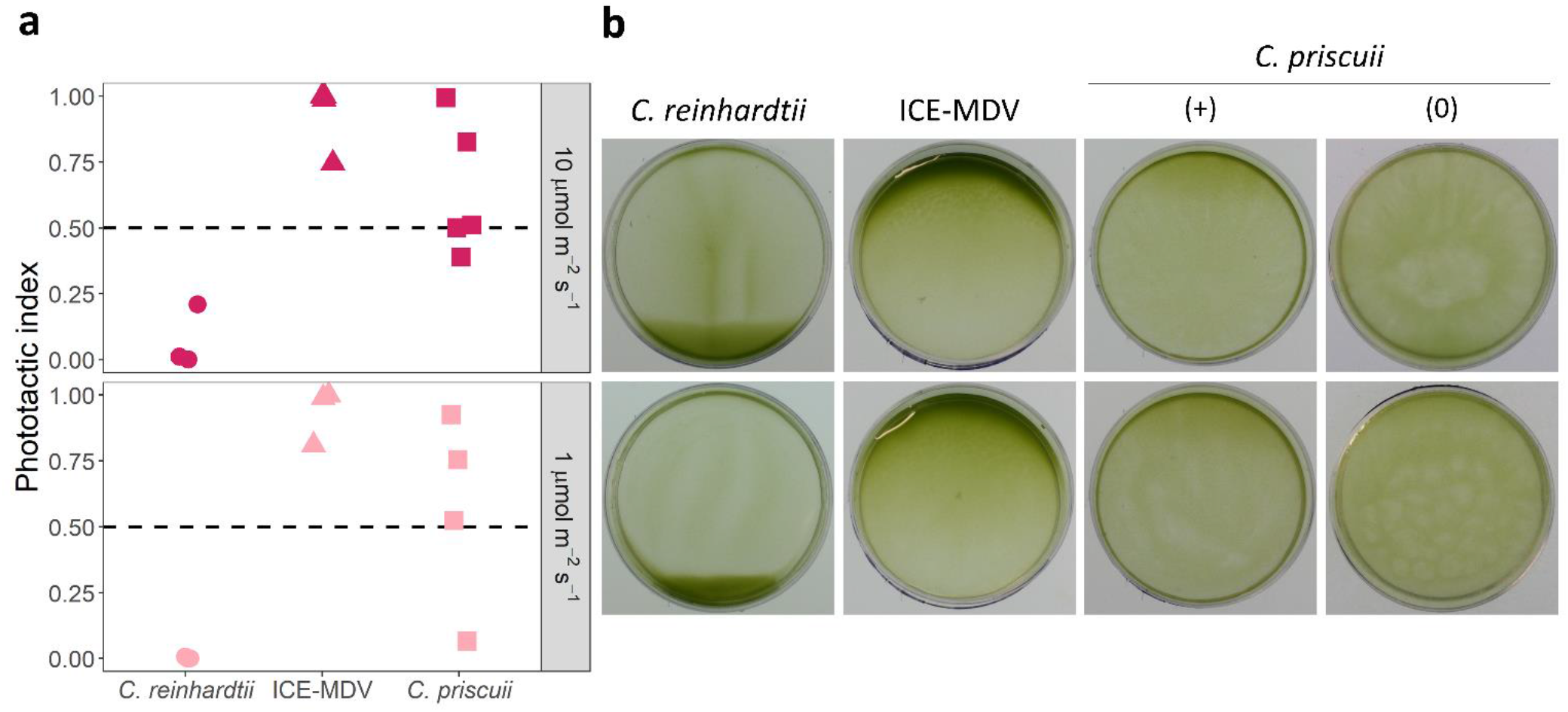
The phototactic response in *C. reinhardtii*, ICE-MDV, and *C. priscuii* determined in a dish motility assay. Cell suspensions were exposed to green light (λ = 525 nm) at two different intensities: 10 μmol m^-2^ s^-1^ (top) and 1 μmol m^-2^ s^-1^ (bottom) and observed after 5 minutes (*C. reinhardtii*) or 15 minutes (ICE-MDV, *C. priscuii*). (**a**) The phototactic index calculated using the pixel density of the images before and after the light treatment. (**b**) Representative images of phototactic movement. *C. priscuii* had a weak or inconsistent phototactic response, and we show a representative image with detectable phototaxis (+) and no phototaxis (0). In all cases, positive phototaxis is indicated by accumulation of cells to the top side of the dish and a phototactic index of 1, negative phototaxis is indicated by accumulation of cells to the bottom side of the dish and a phototactic index of 0, and no phototactic response is seen by uniform dispersal of cells and a phototactic index of 0.5. All experiments were performed as at least 3 biological replicates.

Previous work on *C. reinhardtii* has suggested that phototactic behaviour consists of three steps: 1) photoreception by ChR; 2) a signal transduction pathway that involves Ca^2+^ and reactive oxygen species (ROS); and 3) a change in the beating balance between the two flagella that regulates the phototactic turning (Wakabayashi *et al*., 2021). Thus, the weak or absent phototaxis observed in *C. priscuii* could be a result of a defect in one or more of these steps. To test for flagellar defects, we tested the photoshock response in *C. priscuii*. This ChR-mediated response occurs when algae sense a sudden and strong illumination, which causes a brief stop (<0.5s) and/or a period of backward motion (Baidukova *et al*., 2022). It has been documented that the *C. reinhardtii* mutants ptx1 and lsp1 are not phototactic due to flagellar defects but display clear photoshock response as a result of a functional light-sensing and signalling apparatus (Okita *et al*., 2005). Thus, a lack of a photoshock response would suggest a decreased ability to sense light signals.

A microscope-based photoshock assay revealed that *C. priscuii* has a very weak photoshock response where <10% of cells stopped or reversed their swimming direction when exposed to a very brief (2 ms) bright light flash (Figure 4; Movie S4). In contrast, we observed a robust photoshock response in *C. reinhardtii* (>98% responsive cells) (Figure 4; Movie S5). ICE-MDV displayed a photoshock response, albeit not as strong as the one observed with *C. reinhardtii* (∼65% responsive cells) (Figure 4; Movie S6). These result, and the demonstrated ability of *C. priscuii* to move under non-directional light (Movie S1), suggest that this species has functional flagella but aberrant light-sensing ability.

**Figure 4:**
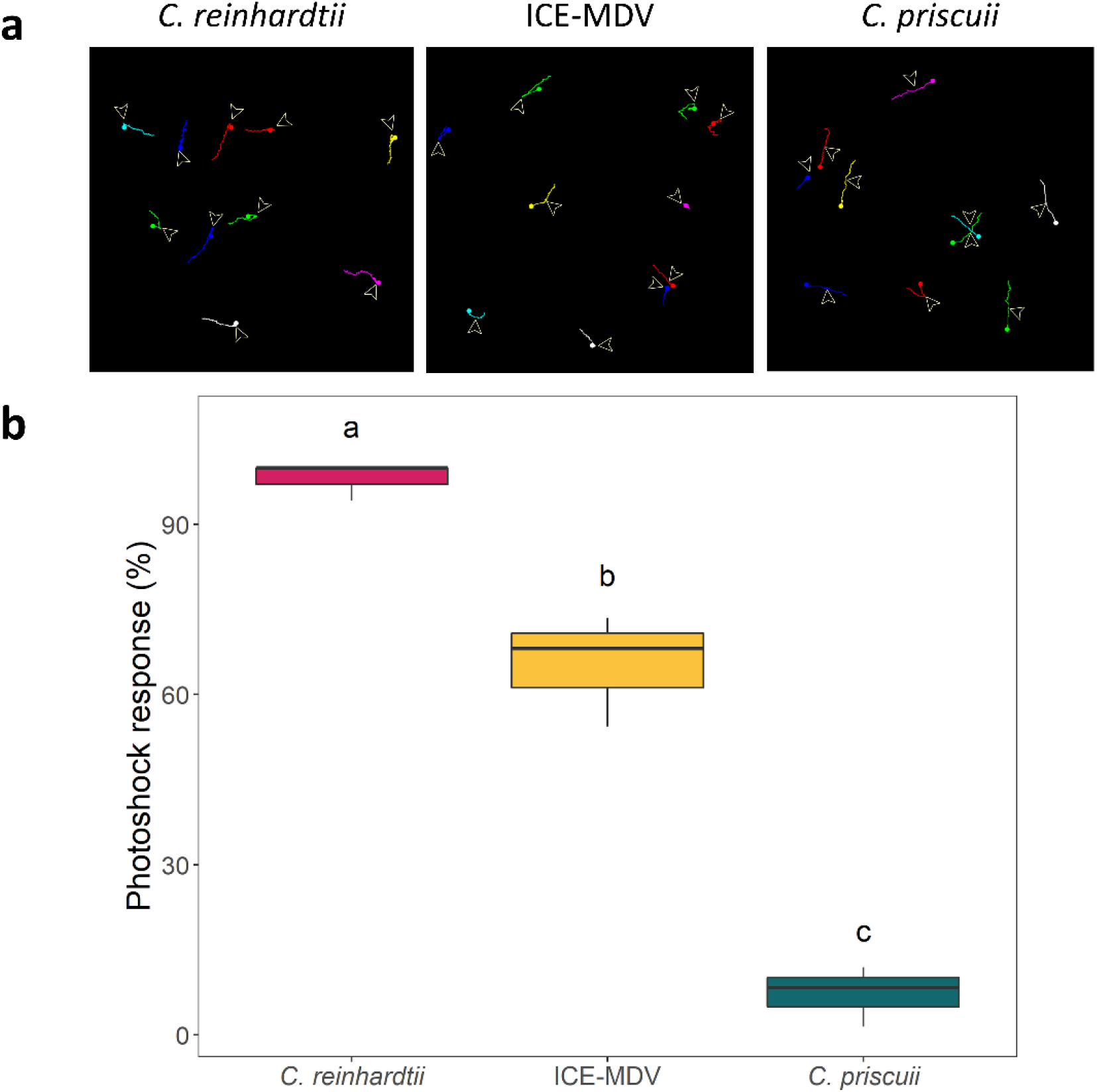
Photoshock response of three Chlamydomonas species. (**a**) Representative 15 second swimming trajectories of *C. reinhardtii*, ICE-MDV, and *C. priscuii* showing cell movement before and after photoshock. The arrow indicates the flash illumination point. Different colors indicate different cell trajectories. (b) Percent of cells that exhibit a photoshock response. Statistical significance was determined by a one-way ANOVA (p = 9.4 x 10^−6^) and Tukey’s post hoc test. Statistically different results are indicated by different letters. (n = 33 cells/species).

### The regulation of phototaxis in psychrophilic algae

The direction of phototaxis is redox regulated in *C. reinhardtii* (Wakabayashi *et al*., 2011), and we tested whether the same is true in its psychrophilic relatives. As shown previously, addition of H_2_O_2_ to the culture media caused positive phototaxis in *C. reinhardtii* exposed to 10 μmol m^-2^ s^-2^ green light, while reactive oxygen species (ROS) scavenging by dimethyl thiourea (DMTU) caused negative phototaxis (Figure 5). The mechanism behind ROS involvement is poorly understood, but it has been postulated that phototactic behavior maintains a moderately reduced state of the cytoplasm and high photosynthetic activity under variable light conditions (Wakabayashi et al., 2011).

**Figure 5:**
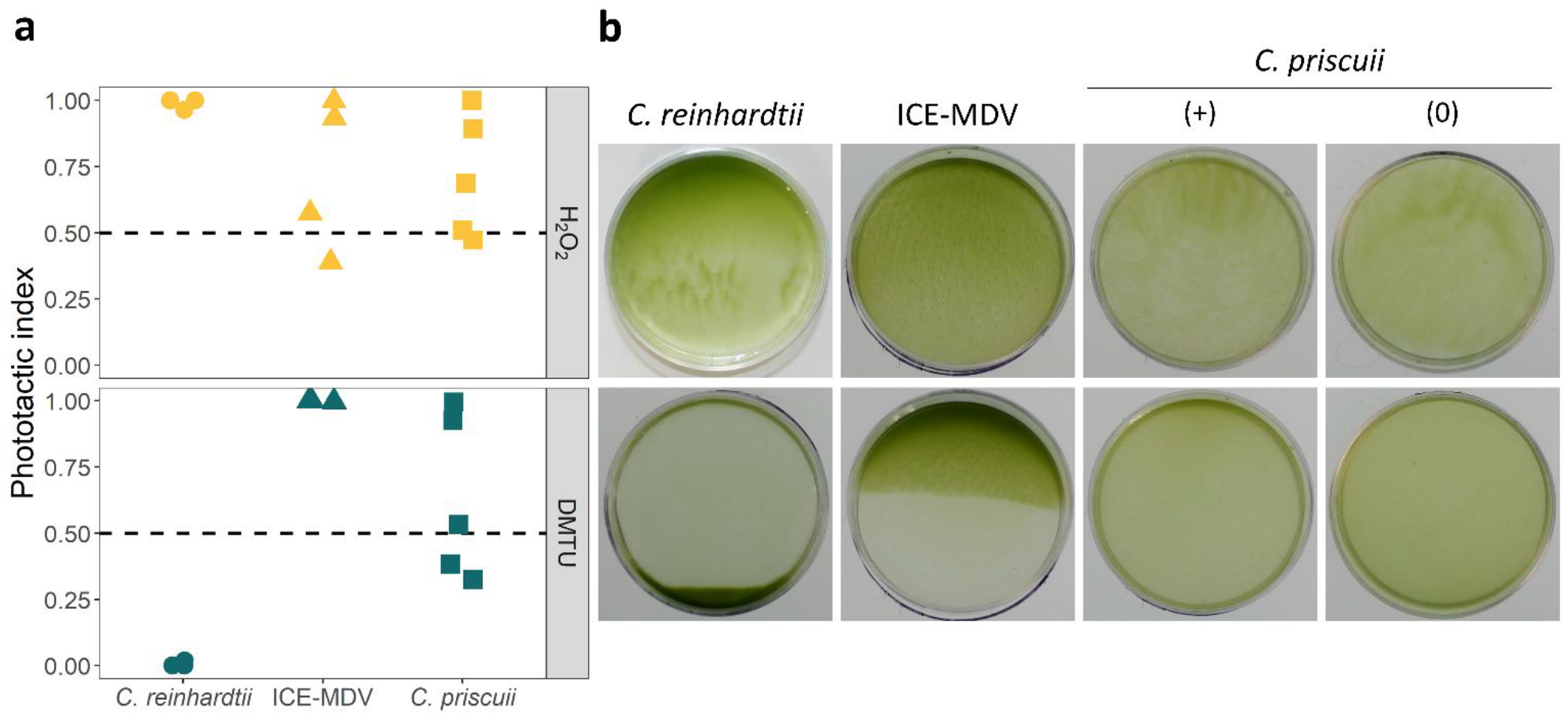
The phototactic response in *C. reinhardtii*, ICE-MDV, and *C. priscuii* treated with H_2_O_2_ (top) and its quencher DMTU (bottom). Cell suspensions were exposed to 10 μmol m^-2^ s^-1^ green light (λ = 525 nm) and observed after 5 minutes (*C. reinhardtii*) or 15 minutes (ICE-MDV, *C. priscii*). (**a**) The phototactic index calculated using the pixel density of the images before and after the light treatment. (**b**) Representative images of phototactic movement. *C. priscuii* had a weak or inconsistent phototactic response, and we show a representative image with detectable phototaxis (+) and no phototaxis (0). In all cases, positive phototaxis is indicated by accumulation of cells to the top side of the dish and a phototactic index of 1, negative phototaxis is indicated by accumulation of cells to the bottom side of the dish and a phototactic index of 0, and no phototactic response is seen by uniform dispersal of cells and a phototactic index of 0.5. All experiments were performed as at least 3 biological replicates.

Our results suggest that psychrophilic algae employ different mechanisms of phototaxis regulation compared to the model *C. reinhardtii*. Addition of H_2_O_2_ had a minimal effect on both Antarctic species (Figure 5). The regulation of phototaxis by H_2_O_2_ appears to be light quality- and quantity-dependent, as all three species displayed weak or absent phototaxis under blue or low intensity green light (Figure S3a). ROS scavenging by DMTU resulted in a very weak positive or no phototactic response in *C. priscuii* but it induced a strong positive phototaxis in ICE-MDV at all light conditions tested here (green and blue light, 1-10 μmol m^-2^ s^-1^) (Figure 5, Figure S3b). Overall, our results suggest that the mechanism behind ROS-dependent regulation of phototaxis is species specific. A recent paper reported that *C. priscuii* has a high capacity for ROS detoxification through constitutive upregulation of the ascorbate pathway (Stahl-Rommel *et al*., 2021).

Next, we examined the role of temperature on the strength of the phototactic response in green algae. Previous work by Pocock *et al*. (2004) observed strong phototaxis in *C. priscuii* cultures in the stationary phase at 25° exposed to high intensity white light, suggesting that the phototactic ability of this alga depends on temperature. To test this, we performed a dish phototactic assay with all three species at suboptimal temperatures (4°C for *C. reinhardtii*; 24°C for *C. priscuii* and ICE-MDV) at 10 μmol m^-2^ s^-1^ of green light, and in the presence of H_2_O_2_ and DMTU. All species were incubated at the experimental temperature up to 2 hours prior to the experiment. In *C. reinhardtii* and ICE-MDV, increased time at suboptimal temperatures lead to a decrease in the phototactic response (Figure S4). We also did not observe a phototactic response in *C. priscuii*, even after 2 hours at 24°C. To test whether the culture growth stage has an effect on motility, we also performed a dish phototaxic assay at the stationary phase for all three species, and we show a decreased motility in *C. reinhardtii* and an absence of motility for ICE-MDV and *C. priscuii* in stationary phase (Figure S5) when compared to actively growing cultures in the mid-log phase (Figure 3). Thus, neither increased temperature nor a different growth stage induced strong phototaxis in *C. priscuii* under environmentally relevant light conditions.

Taken together our results suggest that the Antarctic alga *C. priscuii* has functional flagella but a weak and inconsistent ability to perceive and translate light signals into phototactic motion, regardless of the intensity or quality of light, presence of ROS, temperature or growth stage. We suggest that this is a result of life under extreme shading rather than a common feature of all psychrophilic algae since its close relative ICE-MDV displays a robust ability to move in response to light. Moreover, the high salinity gradient in the deep photic zone of Lake Bonney restricts natural phytoplankton populations from vertical movement in the water column (Lizotte & Priscu, 1994)This fits with earlier studies on native phytoplankton communities in Lake Bonney that demonstrated that shallow water phytoplankton populations had a strong positive phototactic response while deeper water populations had a weak or no phototactic behaviour (Priscu & Neale, 1995).

### Keeping the status quo in Lake Bonney

So, what does this mean for the lifestyle of *C. priscuii* in the deep photic zone of salty Lake Bonney? Our results support the hypothesis that, under physiologically relevant conditions (low blue-green light, high salinity, and low temperature), *C. priscuii* has a limited capability to perceive and translate light signals into rapid motility towards areas of increased light within the water column. This could be due to its small eyespot, reduced photoreceptor repertoire, or altered ROS signalling mechanisms downstream of light perception. A detailed examination of the size and composition of the eyespot, the activity of the psychrophilic ChR photoreceptor, and the signalling pathways that lead to flagellar motility will shed light on phototaxis in shade-adapted green algae.

The extreme conditions in Lake Bonney have undoubtedly shaped the physiology of *C. priscuii* beyond phototaxis. This alga is unable to grow at light intensities >250 μmol m^-2^ s^-1^ (Morgan-Kiss *et al*., 2006), and lacks short-term photoacclimation response for balancing light energy distribution between the photosystems via state transitions, which is well conserved in many other algal species (Morgan-Kiss *et al*., 2002; Szyszka *et al*., 2007). Instead, it accumulates a unique PSI-cyt *b*_*6*_*f* photosynthetic supercomplex (Szyszka-Mroz *et al*., 2015), which supports constitutively active cyclic electron flow (CEF) around PSI for energy homeostasis and photoprotection (Kalra *et al*., 2020). These features have been associated with a rewired primary carbon metabolism leading to constitutively high levels of stress-related compounds (e.g., sucrose, proline, and antioxidants) that support robust growth under extreme conditions (Kalra *et al*., 2020; Stahl-Rommel *et al*., 2021; Cvetkovska *et al*., 2022). It is likely that the combined pressures of low temperatures, hypersalinity, and extreme shading drive these adjustments in its physiology, including a reduced phototactic response. Examination of the evolutionary history of *C. priscuii* and its arrival in Lake Bonney would further inform on its unique physiology and behaviour.

Lake Bonney is highly stratified and nutrient limited, particularly for phosphorus (Priscu, 1995). Priscu and Neale (1995) proposed that nutrient limitation, rather than light, may drive the tactic behavior of phytoflagellates residing in the deep photic zone. They postulated that motile phytoflagellates have a competitive advantage over nonmotile algae for maintaining their position in the photic zone at discrete depths in response to trade-offs between light utilization efficiency and nutrient availability (Priscu & Neale, 1995). Indeed, in nutrient bioassay experiments, Lake Bonney *Chlamydomonas* spp. were highly competitive under phosphorus-supplemented treatments (Teufel *et al*., 2016). Recent models suggest that polar environments will be strongly affected by climate change and ice-free conditions may disrupt the lake stratification, nutrient and light availaibility (Obryk *et al*., 2019). For instance, climate-driven high stream flow years carry significant levels of nitrogen and phosphorus to the nutrient-poor waters of Lake Bonney (Gooseff *et al*., 2017). How these complex climate-driven processes disrupting the ‘status quo’ of Lake Bonney will affect the endemic and highly specialized psychrophiles, such as *C. priscuii* and ICE-MDV, is currently not known. Studying the physiology and environmental responses of these unique organisms is important for predicting their survival in the face of global climate change.

## Data availability

The genomic data that support the findings of this study are available in Phytozome (https://phytozome-next.jgi.doe.gov/) and NCBI GenBank (https://www.ncbi.nlm.nih.gov/genbank/). All accession numbers for the sequences are available within the Supplementary Data (Supplementary Table S1). All other data that support the findings of this study are available from the corresponding author upon reasonable request.

## Supplementary material

**Table S1**: Photoreceptor genes in *Chlamydomonas priscuii* and related Chlamydomonadales. The data was obtained from GenBank for *C. priscuii* (PRJNA547753) and *Chlamydomonas* sp. ICE-L (PRJNA636631). All other algal genomes were obtained from PhycoCosm. The table contains the accession numbers for all identified genes.

**Figure S1:** Multiple sequence alignments of channelrhodopsins from different Chlamydomonadalean species. Residues shown to be phosphorylated in *C. reinhardtii* (Böhm *et al*., 2019) are highlighted with a yellow square and those that are also present in *C. priscuii* are labeled with a red asterisk. The retinal binding sites are labeled with a blue square (Kato *et al*., 2012).

**Figure S2**: The phototactic response in *C. reinhardtii*, ICE-MDV, and *C. priscuii* exposed to blue light (λ = 420 nm) at two different intensities: 10 μmol m^-2^ s^-1^ (top) and 1 μmol m^-2^ s^-1^ (bottom). (**a**) Phototactic index (**b**) Representative images from a dish motility assay. All experiments were performed in at least 3 biological replicates.

**Figure S3:** Phototactic response represented as a phototactic index for *C. reinhardtii*, ICE-MDV, and *C. priscuii* in the presence of (**a**) H_2_O_2_ and (**b**) DMTU under blue light (10 μmol m^-2^ s^-1^, 1 μmol m^-2^ s^-1^) and green light (1 μmol m^-2^ s^-1^). All experiments were performed in at least 3 biological replicates.

**Figure S4:** Phototactic response in *C. reinhardtii*, ICE-MDV, and *C. priscuii* after incubation at suboptimal temperatures. Cell suspensions were incubated at 4°C (*C. reinhardtii*) or 24°C (ICE-MDV and *C. priscuii)* for 30 minutes or 2 hours prior to phototaxis dish assay. Cell suspensions were exposed to 10 μmol m^-2^ s^-1^ green light with and without the addition of H_2_O_2_ or DMTU (**a**) The phototactic index was calculated using the pixel density images (**b**) Representative images. All experiments were performed in at least 3 biological replicates.

**Figure S5**: Phototactic response in *C. reinhardtii*, ICE-MDV, and *C. priscuii* at stationary phase exposed to 10 μmol m^-2^ s^-1^ green light with and without the addition of H_2_O_2_ or DMTU. (**a**) The phototactic index was calculated using the pixel density images (**b**). Representative images. All experiments were performed in at least 3 biological replicates.

**Movie S1**: Motility in *C. priscuii* under non-directional white light. Scale bar = 100 μm. (**Submitted as an AVI file**)

**Movie S2**: Motility in *C. reinhardtii* under non-directional white light. Scale bar = 100 μm. (**Submitted as an AVI file**)

**Movie S3**: Motility in ICE-MDV under non-directional white light. Scale bar = 100 μm. (**Submitted as an AVI file**)

**Movie S4**: Photoshock response of in *C. priscuii* under non-directional 5 μmol m^-2^ s^-1^ red light followed by exposure to a white light flash for 2 ms. Scale bar = 100 μm. (**Submitted as an AVI file**)

**Movie S5**: Photoshock response of in *C. reinhardtii* under non-directional 5 μmol m^-2^ s^-1^ red light followed by exposure to a white light flash for 2 ms. Scale bar = 100 μm. (**Submitted as an AVI file**)

**Movie S6**: Photoshock response of in ICE-MDV under non-directional 5 μmol m^-2^ s^-1^ red light followed by exposure to a white light flash for 2 ms. Scale bar = 100 μm. (**Submitted as an AVI file**)

## References

Baidukova O, Oppermann J, Kelterborn S, Fernandez Lahore RG, Schumacher D, Evers H, Kamrani YY, Hegemann P. 2022. Gating and ion selectivity of Channelrhodopsins are critical for photo-activated orientation of Chlamydomonas as shown by in vivo point mutation. Nature Communications 13: 7253.

Beel B, Prager K, Spexard M, Sasso S, Weiss D, Müller N, Heinnickel M, Dewez D, Ikoma D, Grossman AR, et al. 2012. A Flavin Binding Cryptochrome Photoreceptor Responds to Both Blue and Red Light in Chlamydomonas reinhardtii. The Plant Cell 24: 2992–3008.

Berthold P, Tsunoda SP, Ernst OP, Mages W, Gradmann D, Hegemann P. 2008. Channelrhodopsin-1 initiates phototaxis and photophobic responses in Chlamydomonas by immediate light-induced depolarization. The Plant Cell 20: 1665–1677.

Bielewicz S, Bell E, Kong W, Friedberg I, Priscu JC, Morgan-Kiss RM. 2011. Protist diversity in a permanently ice-covered Antarctic Lake during the polar night transition. The ISME Journal 5: 1559–1564.

Böhm M, Boness D, Fantisch E, Erhard H, Frauenholz J, Kowalzyk Z, Marcinkowski N, Kateriya S, Hegemann P, Kreimer G. 2019. Channelrhodopsin-1 Phosphorylation Changes with Phototactic Behavior and Responds to Physiological Stimuli in Chlamydomonas. The Plant Cell 31: 886–910.

Böhm M, Kreimer G. 2020. Orient in the World with a Single Eye: The Green Algal Eyespot and Phototaxis. In: Cánovas FM, Lüttge U, Risueño M-C, Pretzsch H, eds. Progress in Botany. Progress in Botany Vol. 82. Cham: Springer International Publishing, 259–304.

Craig RJ, Gallaher SD, Shu S, Salomé P, Jenkins JW, Blaby-Haas CE, Purvine SO, O’Donnell S, Barry K, Grimwood J, et al. 2022. The Chlamydomonas Genome Project, version 6: reference assemblies for mating type plus and minus strains reveal extensive structural mutation in the laboratory. The Plant Cell: koac347.

Craig RJ, Hasan AR, Ness RW, Keightley PD. 2021. Comparative genomics of Chlamydomonas. The Plant Cell 33: 1016–1041.

Cvetkovska M, Zhang X, Vakulenko G, Benzaquen S, Szyszka-Mroz B, Malczewski N, Smith DR, Hüner NPA. 2022. A constitutive stress response is a result of low temperature growth in the Antarctic green alga Chlamydomonas sp. UWO241. Plant, Cell & Environment 45: 156–177.

Gavelis GS, Keeling PJ, Leander BS. 2017. How exaptations facilitated photosensory evolution: Seeing the light by accident. BioEssays 39: 1600266.

Gooseff MN, Barrett JE, Adams BJ, Doran PT, Fountain AG, Lyons WB, McKnight DM, Priscu JC, Sokol ER, Takacs-Vesbach C, et al. 2017. Decadal ecosystem response to an anomalous melt season in a polar desert in Antarctica. Nature Ecology & Evolution 1: 1334–1338.

Greiner A, Kelterborn S, Evers H, Kreimer G, Sizova I, Hegemann P. 2017. Targeting of Photoreceptor Genes in Chlamydomonas reinhardtii via Zinc-Finger Nucleases and CRISPR/Cas9. The Plant Cell 29: 2498–2518.

Grigoriev IV, Hayes RD, Calhoun S, Kamel B, Wang A, Ahrendt S, Dusheyko S, Nikitin R, Mondo SJ, Salamov A, et al. 2021. PhycoCosm, a comparative algal genomics resource. Nucleic Acids Research 49: D1004–D1011.

Hanschen ER, Marriage TN, Ferris PJ, Hamaji T, Toyoda A, Fujiyama A, Neme R, Noguchi H, Minakuchi Y, Suzuki M, et al. 2016. The Gonium pectorale genome demonstrates co-option of cell cycle regulation during the evolution of multicellularity. Nature Communications 7: 11370.

Hirooka S, Hirose Y, Kanesaki Y, Higuchi S, Fujiwara T, Onuma R, Era A, Ohbayashi R, Uzuka A, Nozaki H, et al. 2017. Acidophilic green algal genome provides insights into adaptation to an acidic environment. Proceedings of the National Academy of Sciences. 114: E8304–E8313

Jékely G. 2009. Evolution of phototaxis. Philosophical Transactions of the Royal Society B: Biological Sciences 364: 2795–2808.

Kalra I, Wang X, Cvetkovska M, Jeong J, McHargue W, Zhang R, Hüner N, Yuan JS, Morgan-Kiss R. 2020. Chlamydomonas sp. UWO 241 Exhibits High Cyclic Electron Flow and Rewired Metabolism under High Salinity. Plant Physiology 183: 588–601.

Kato HE, Zhang F, Yizhar O, Ramakrishnan C, Nishizawa T, Hirata K, Ito J, Aita Y, Tsukazaki T, Hayashi S, et al. 2012. Crystal structure of the channelrhodopsin light-gated cation channel. Nature 482: 369–374.

Kianianmomeni A, Hallmann A. 2014. Algal photoreceptors: in vivo functions and potential applications. Planta 239: 1–26.

Kong W, Ream DC, Priscu JC, Morgan-Kiss RM. 2012. Diversity and Expression of RubisCO Genes in a Perennially Ice-Covered Antarctic Lake during the Polar Night Transition. Applied and Environmental Microbiology 78: 4358–4366.

Li W, Podar M, Morgan-Kiss RM. 2016. Ultrastructural and Single-Cell-Level Characterization Reveals Metabolic Versatility in a Microbial Eukaryote Community from an Ice-Covered Antarctic Lake. Applied and Environmental Microbiology.

Li F-W, Rothfels CJ, Melkonian M, Villarreal JC, Stevenson DW, Graham SW, Wong GK-S, Mathews S, Pryer KM. 2015. The origin and evolution of phototropins. Frontiers in Plant Science 6: 637.

Lizotte MP, Sharp TR, Priscu JC. 1996. Phytoplankton dynamics in the stratified water column of Lake Bonney, Antarctica. Polar Biology 16: 155–162.

Lu S, Wang J, Chitsaz F, Derbyshire MK, Geer RC, Gonzales NR, Gwadz M, Hurwitz DI, Marchler GH, Song JS, et al. 2020. CDD/SPARCLE: the conserved domain database in 2020. Nucleic Acids Research 48: D265–D268.

Luck M, Hegemann P. 2017. The two parallel photocycles of the Chlamydomonas sensory photoreceptor histidine kinase rhodopsin 1. Journal of Plant Physiology 217: 77–84.

Luck M, Mathes T, Bruun S, Fudim R, Hagedorn R, Nguyen TMT, Kateriya S, Kennis JTM, Hildebrandt P, Hegemann P. 2012. A Photochromic Histidine Kinase Rhodopsin (HKR1) That Is Bimodally Switched by Ultraviolet and Blue Light. Journal of Biological Chemistry 287: 40083–40090.

Merchant SS, Prochnik SE, Vallon O, Harris EH, Karpowicz SJ, Witman GB, Terry A, Salamov A, Fritz-Laylin LK, Maréchal-Drouard L, et al. 2007. The Chlamydomonas genome reveals the evolution of key animal and plant functions. Science (New York, N.Y.) 318: 245–250.

Mistry J, Chuguransky S, Williams L, Qureshi M, Salazar GA, Sonnhammer ELL, Tosatto SCE, Paladin L, Raj S, Richardson LJ, et al. 2021. Pfam: The protein families database in 2021. Nucleic Acids Research 49: D412–D419.

Morgan-Kiss R, Ivanov AG, Williams J, Mobashsher Khan, Huner NPA. 2002. Differential thermal effects on the energy distribution between photosystem II and photosystem I in thylakoid membranes of a psychrophilic and a mesophilic alga. Biochimica et Biophysica Acta (BBA) - Biomembranes 1561: 251–265.

Morgan-Kiss RM, Priscu JC, Pocock T, Gudynaite-Savitch L, Huner NPA. 2006. Adaptation and Acclimation of Photosynthetic Microorganisms to Permanently Cold Environments. Microbiology and Molecular Biology Reviews 70: 222–252.

Müller N, Wenzel S, Zou Y, Künzel S, Sasso S, Weiß D, Prager K, Grossman A, Kottke T, Mittag M. 2017. A Plant Cryptochrome Controls Key Features of the Chlamydomonas Circadian Clock and Its Life Cycle. Plant Physiology 174: 185–201.

Nakasone Y, Ohshima M, Okajima K, Tokutomi S, Terazima M. 2018. Photoreaction Dynamics of LOV1 and LOV2 of Phototropin from Chlamydomonas reinhardtii. The Journal of Physical Chemistry B 122: 1801–1815.

Neale PJ, Priscu JC. 1995. The Photosynthetic Apparatus of Phytoplankton from a Perennially Ice-Covered Antarctic Lake: Acclimation to an Extreme Shade Environment. Plant and Cell Physiology 36: 253–263.

Obryk MK, Doran PT, Priscu JC. 2019. Prediction of Ice–Free Conditions for a Perennially Ice–Covered Antarctic Lake. Journal of Geophysical Research: Earth Surface 124: 686–694.

Okita N, Isogai N, Hirono M, Kamiya R, Yoshimura K. 2005. Phototactic activity in Chlamydomonas ‘non-phototactic’ mutants deficient in Ca2+-dependent control of flagellar dominance or in inner-arm dynein. Journal of Cell Science 118: 529–537.

Patriarche JD, Priscu JC, Takacs-Vesbach C, Winslow L, Myers KF, Buelow H, Morgan-Kiss RM, Doran PT. 2021. Year-Round and Long-Term Phytoplankton Dynamics in Lake Bonney, a Permanently Ice-Covered Antarctic Lake. Journal of Geophysical Research: Biogeosciences 126: e2020JG005925.

Petroutsos D, Tokutsu R, Maruyama S, Flori S, Greiner A, Magneschi L, Cusant L, Kottke T, Mittag M, Hegemann P, et al. 2016. A blue-light photoreceptor mediates the feedback regulation of photosynthesis. Nature 537: 563–566.

Pocock T, Lachance M-A, Pröschold T, Priscu JC, Kim SS, Huner NPA. 2004. Identification of a psychrophilic green alga from Lake Bonney Antarctica: Chlamydomonas raudensis Ettl. (UWO241) Chlorophyceae. Journal of Phycology 40: 1138–1148.

Polle JEW, Barry K, Cushman J, Schmutz J, Tran D, Hathwaik LT, Yim WC, Jenkins J, McKie-Krisberg Z, Prochnik S, et al. 2017. Draft Nuclear Genome Sequence of the Halophilic and Beta-Carotene-Accumulating Green Alga Dunaliella salina Strain CCAP19/18. Genome Announcements 5.

Possmayer M, Berardi G, Beall BFN, Trick CG, Hüner NPA, Maxwell DP. 2011. Plasticity of the psychrophilic green alga Chlamydomonas raudensis (UWO 241) (Chlorophyta) to supraoptimal temperature stress: Heat stress in a psychrophilic green alga.. Journal of Phycology 47: 1098–1109.

Possmayer M, Gupta RK, Szyszka–Mroz B, Maxwell DP, Lachance M, Hüner NPA, Smith DR. 2016. Resolving the phylogenetic relationship between Chlamydomonas sp. UWO 241 and Chlamydomonas raudensis sag 49.72 (Chlorophyceae) with nuclear and plastid DNA sequences (M Schroda, Ed.). Journal of Phycology 52: 305–310.

Priscu JC. 1995. Phytoplankton nutrient deficiency in lakes of the McMurdo dry valleys, Antarctica. Freshwater Biology 34: 215–227.

Priscu J, Neale P. 1995. Phototactic response of phytoplankton forming discrete layers within the water column of Lake Bonney, Antarctica. Antarctic Journal of the US 30: 301–303.

Prochnik SE, Umen J, Nedelcu AM, Hallmann A, Miller SM, Nishii I, Ferris P, Kuo A, Mitros T, Fritz-Laylin LK, et al. 2010. Genomic analysis of organismal complexity in the multicellular green alga Volvox carteri. Science (New York, N.Y.) 329: 223–226.

Qian W, Zhang J. 2014. Genomic evidence for adaptation by gene duplication. Genome Research 24: 1356–1362.

Raymond JA, Morgan-Kiss R. 2013. Separate Origins of Ice-Binding Proteins in Antarctic Chlamydomonas Species. PLOS ONE 8: e59186.

Rredhi A, Petersen J, Schubert M, Li W, Oldemeyer S, Li W, Westermann M, Wagner V, Kottke T, Mittag M. 2021. DASH cryptochrome 1, a UV–A receptor, balances the photosynthetic machinery of Chlamydomonas reinhardtii. New Phytologist 232: 610–624.

Schneider CA, Rasband WS, Eliceiri KW. 2012. NIH Image to ImageJ: 25 years of image analysis. Nature Methods 9: 671–675.

Sievers F, Wilm A, Dineen D, Gibson TJ, Karplus K, Li W, Lopez R, McWilliam H, Remmert M, Söding J, et al. 2011. Fast, scalable generation of high-quality protein multiple sequence alignments using Clustal Omega. Molecular Systems Biology 7: 539.

Stahl-Rommel S, Kalra I, D’Silva S, Hahn MM, Popson D, Cvetkovska M, Morgan-Kiss RM. 2021. Cyclic electron flow (CEF) and ascorbate pathway activity provide constitutive photoprotection for the photopsychrophile, Chlamydomonas sp. UWO 241 (renamed Chlamydomonas priscuii). Photosynthesis Research 151: 235–250

Szyszka B, Ivanov AG, Hüner NPA. 2007. Psychrophily is associated with differential energy partitioning, photosystem stoichiometry and polypeptide phosphorylation in Chlamydomonas raudensis. Biochimica et Biophysica Acta (BBA) - Bioenergetics 1767: 789–800.

Szyszka-Mroz B, Pittock P, Ivanov AG, Lajoie G, Hüner NPA. 2015. The Antarctic Psychrophile Chlamydomonas sp. UWO 241 Preferentially Phosphorylates a Photosystem I-Cytochrome b6/f Supercomplex. Plant Physiology 169: 717–736.

Teufel AG, Li W, Kiss AJ, Morgan-Kiss RM. 2016. Impact of nitrogen and phosphorus on phytoplankton production and bacterial community structure in two stratified Antarctic lakes: a bioassay approach. Polar Biology 5: 1007–1022.

Trippens J, Greiner A, Schellwat J, Neukam M, Rottmann T, Lu Y, Kateriya S, Hegemann P, Kreimer G. 2012. Phototropin Influence on Eyespot Development and Regulation of Phototactic Behavior in Chlamydomonas reinhardtii. The Plant Cell 24: 4687–4702.

Ueki N, Ide T, Mochiji S, Kobayashi Y, Tokutsu R, Ohnishi N, Yamaguchi K, Shigenobu S, Tanaka K, Minagawa J, et al. 2016. Eyespot-dependent determination of the phototactic sign in Chlamydomonas reinhardtii. Proceedings of the National Academy of Sciences 113: 5299–5304.

Ueki N, Isu A, Kyuji A, Asahina Y, So S, Takahashi R, Hisabori T, Wakabayashi K-I. 2022. Observation of Photobehavior in Chlamydomonas reinhardtii. Journal of Visualized Experiments: JoVE.

Ueki N, Wakabayashi K. 2017. Phototaxis Assay for Chlamydomonas reinhardtii. BIO-PROTOCOL 7.

Wakabayashi K, Isu A, Ueki N. 2021. Channelrhodopsin-Dependent Photo-Behavioral Responses in the Unicellular Green Alga Chlamydomonas reinhardtii. In: Yawo H, Kandori H, Koizumi A, Kageyama R, eds. Advances in Experimental Medicine and Biology. Optogenetics: Light-Sensing Proteins and Their Applications in Neuroscience and Beyond. Singapore: Springer, 21–33.

Wakabayashi K., Misawa Y, Mochiji S, Kamiya R. 2011. Reduction-oxidation poise regulates the sign of phototaxis in Chlamydomonas reinhardtii. Proceedings of the National Academy of Sciences 108: 11280–11284.

Zhang X, Cvetkovska M, Morgan-Kiss R, Hüner NPA, Smith DR. 2021. Draft genome sequence of the Antarctic green alga Chlamydomonas sp. UWO241. iScience 24: 102084.

Zhang Z, Qu C, Zhang K, He Y, Zhao X, Yang L, Zheng Z, Ma X, Wang X, Wang W, et al. 2020. Adaptation to Extreme Antarctic Environments Revealed by the Genome of a Sea Ice Green Alga. Current Biology 30: 3330-3341.e7.

