## Supplementary figures and images for "Aberrant light sensing and motility in the green alga *Chlamydomonas priscuii* from the ice-covered Antarctic Lake Bonney"

### Figure S1

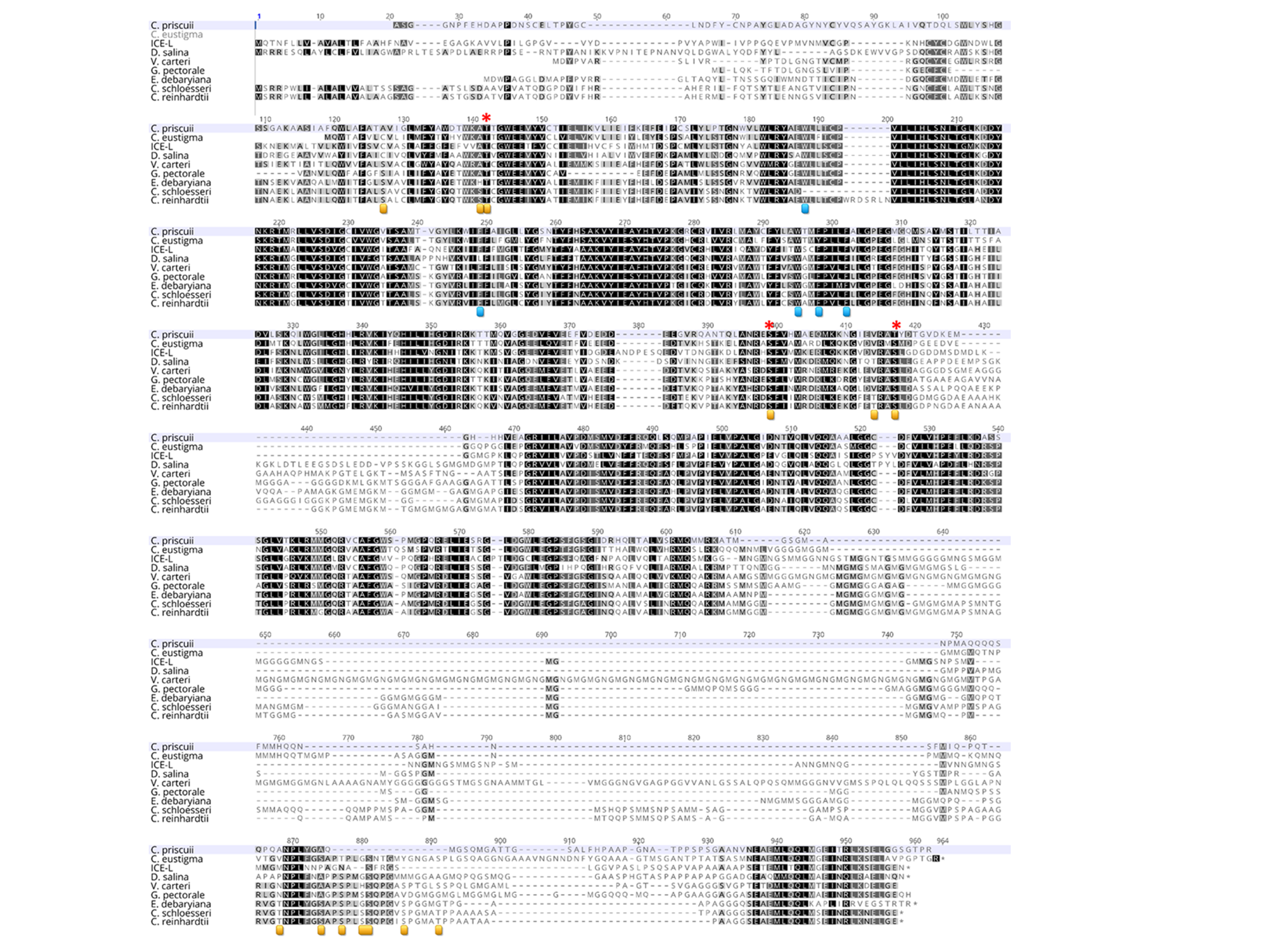

### Figure S2

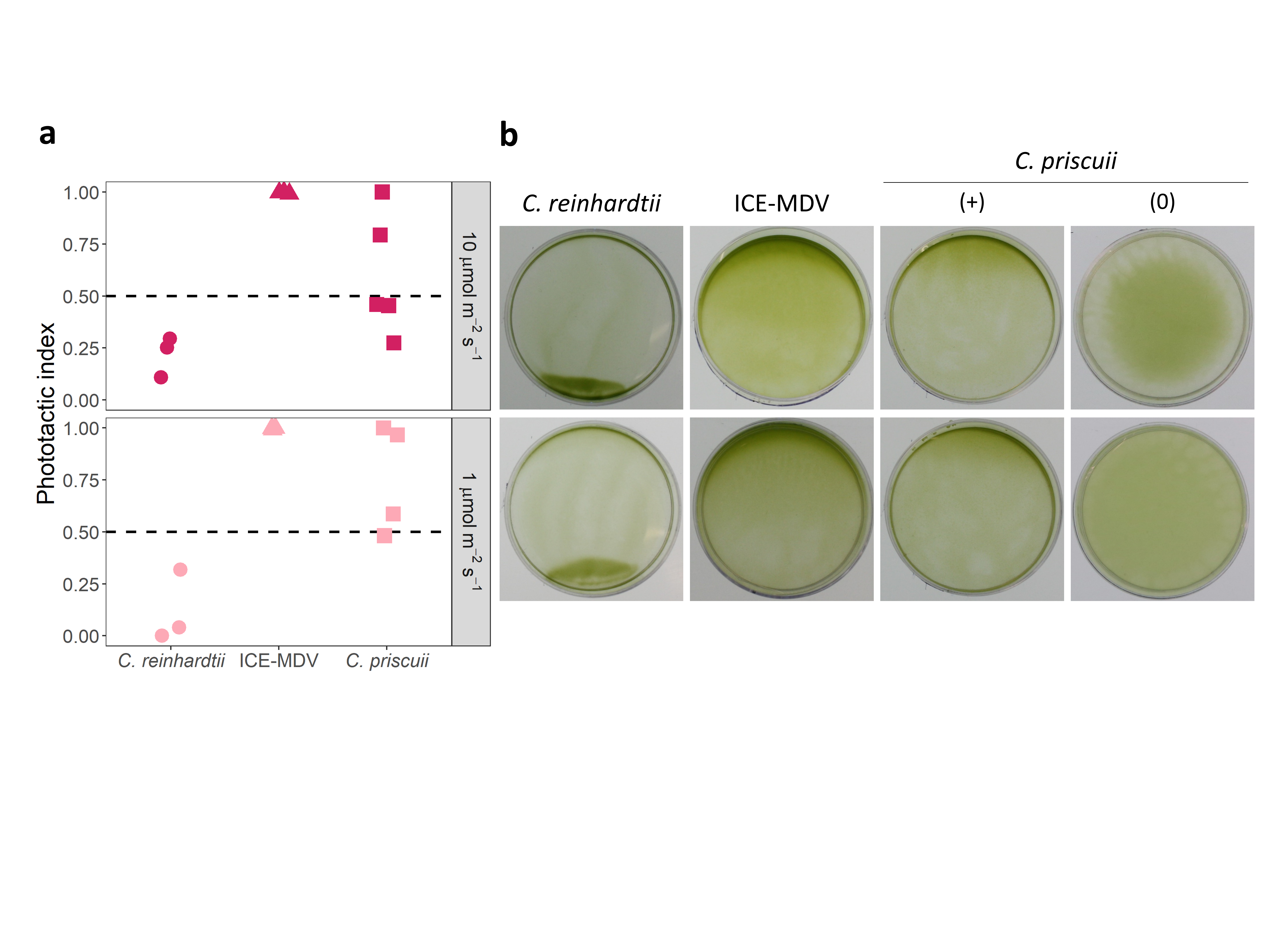

### Figure S3

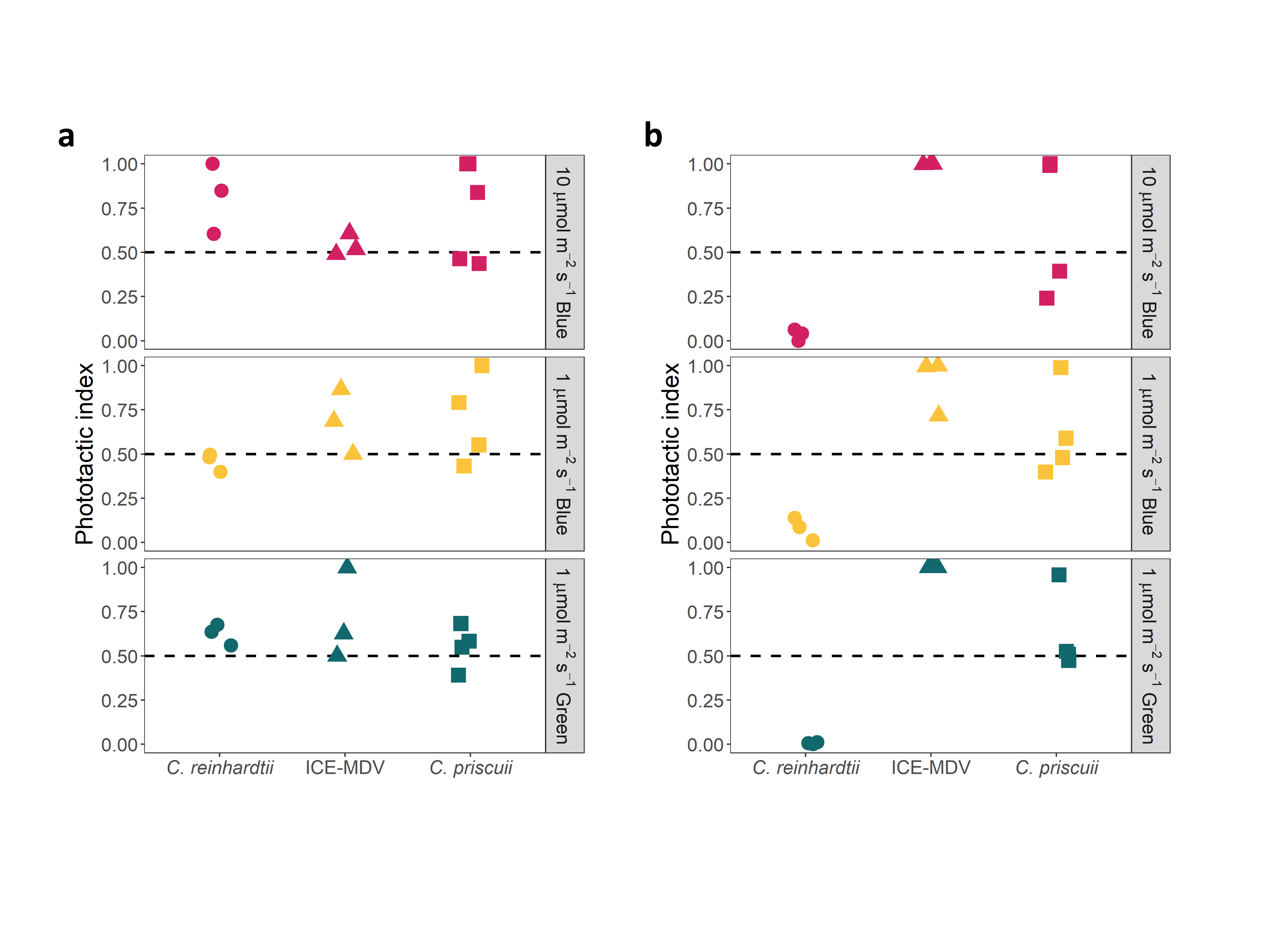

### Figure S4

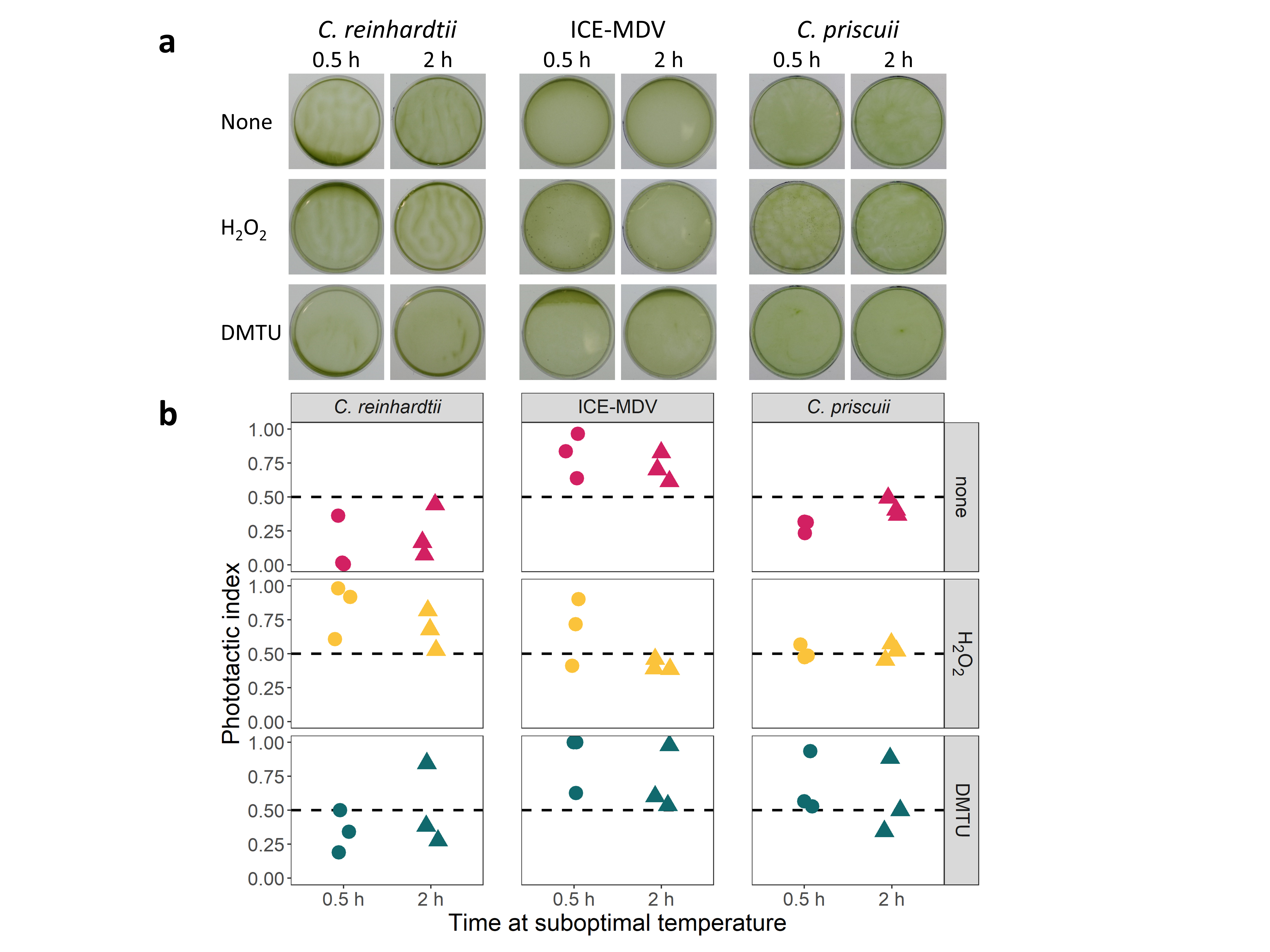

### Figure S5

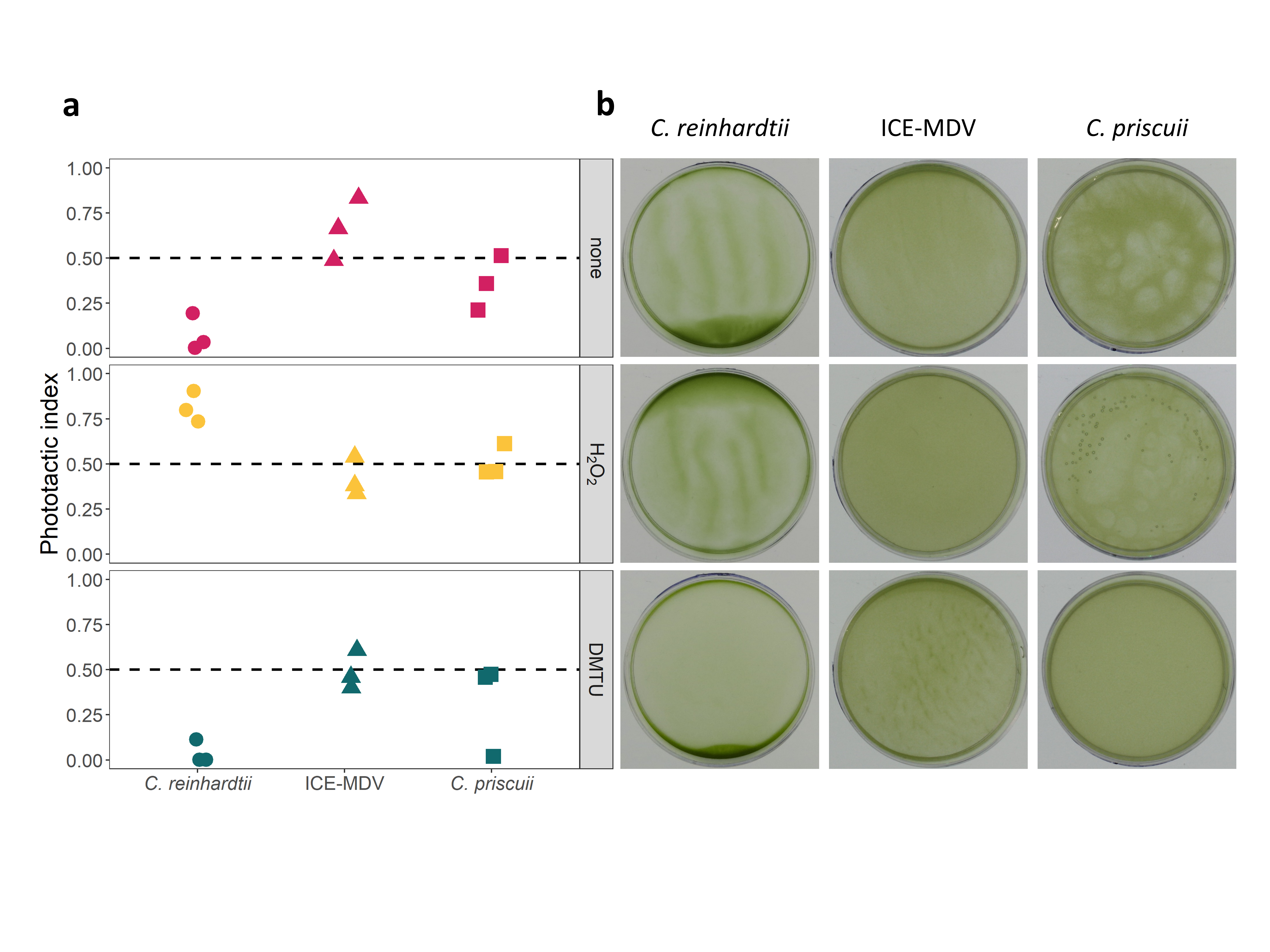
